## Supplementary Fig 1 to 5 for "Enhanced niche colonisation and competition during bacterial adaptation to a fungus"

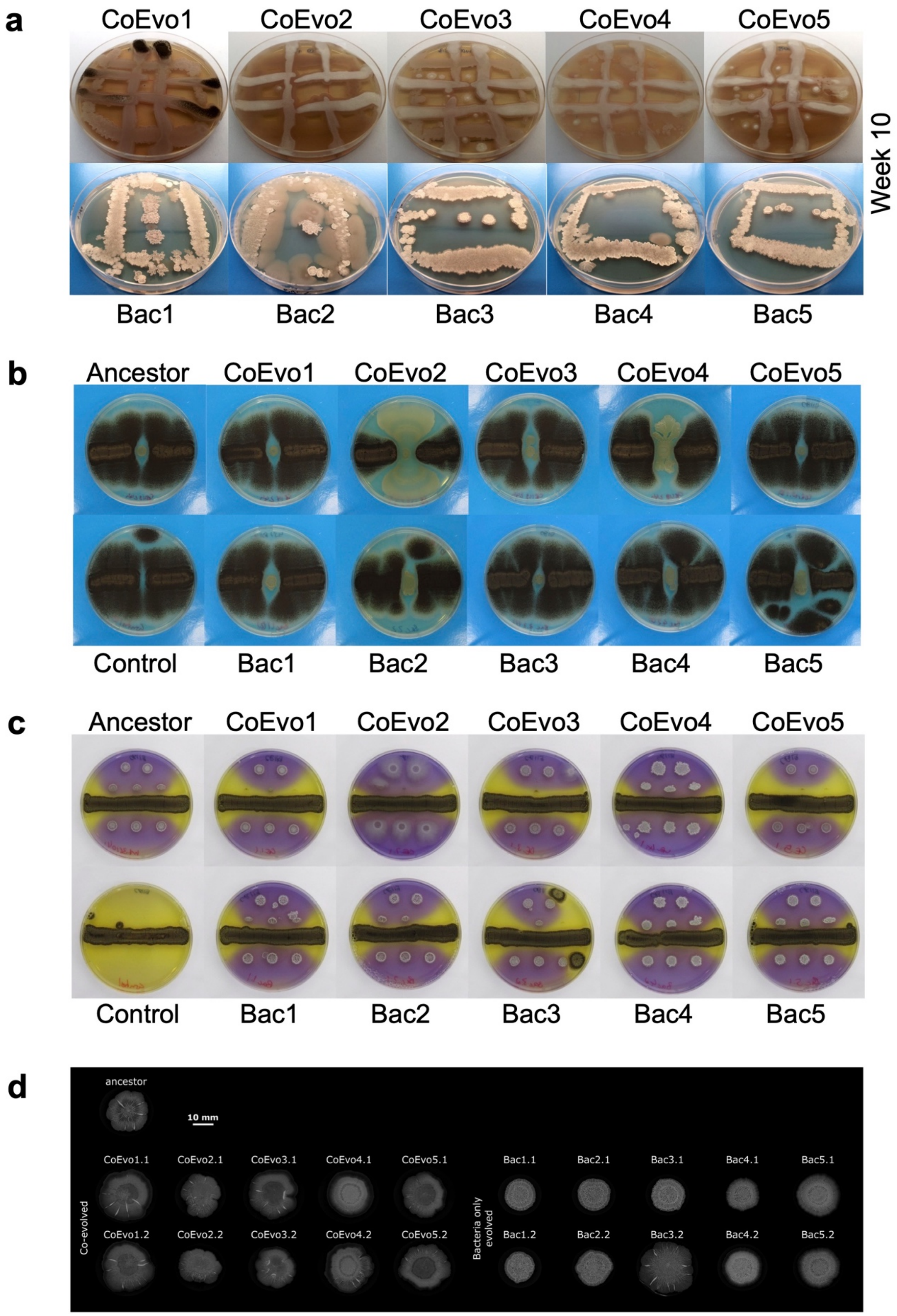


**Fig. S1 | *B. subtilis* adaptation to the presence of *A. niger*. a,** Experimental evolution plates at the 10th transfer, top panels showing *B. subtilis* evolved in the presence of *A. niger*, and lower panels showing bacteria only cultivations. **b,** Bacterial growth spotted between two lines of fungal streaks. **c,** Lower panels showing the pH of the medium using Bromocresol Purple, where bacterial cultures were spotted next to a continuous fungal spore streak. Purple and yellow colours indicate pH >6.8 and pH <5.2, respectively. In **b** and **c**, the plate size = 9 cm, CoEvo refers to co-culture evolved isolates, Bac denotes bacteria only evolved isolated. **d,** Biofilm colonies of evolved isolates on MSgg agar medium. Scale bar = 10 mm.

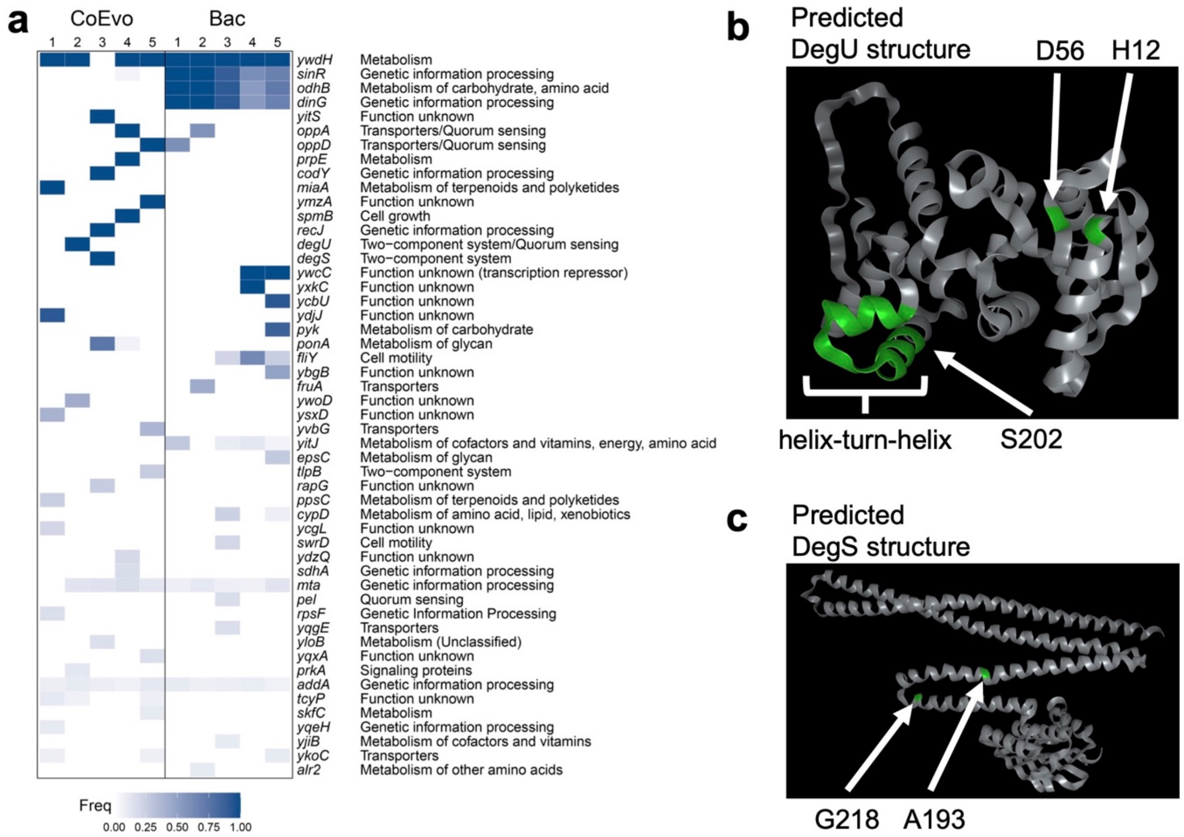


**Fig. S2 | Genetic characterisation of *B. subtilis* adaptation to *A. niger*. a,** Detected mutations in CoEvo and Bac populations. **b,** Predicted DegU structure based on AlphaFold (https://alphafold.ebi.ac.uk/entry/P13800). H12 and D56 amino acids are highlighted that were previously described to be involved in phosphorylation state of DegU. SNP in CoEvo2, S202 is also highlighted. **c,** Predicted DegS structure based on AlphaFold (https://alphafold.ebi.ac.uk/entry/P13799). G218 amino acid is highlighted that were previously described to be involved in phosphor-activity of DegS. SNP in CoEvo3, A193 is also highlighted.

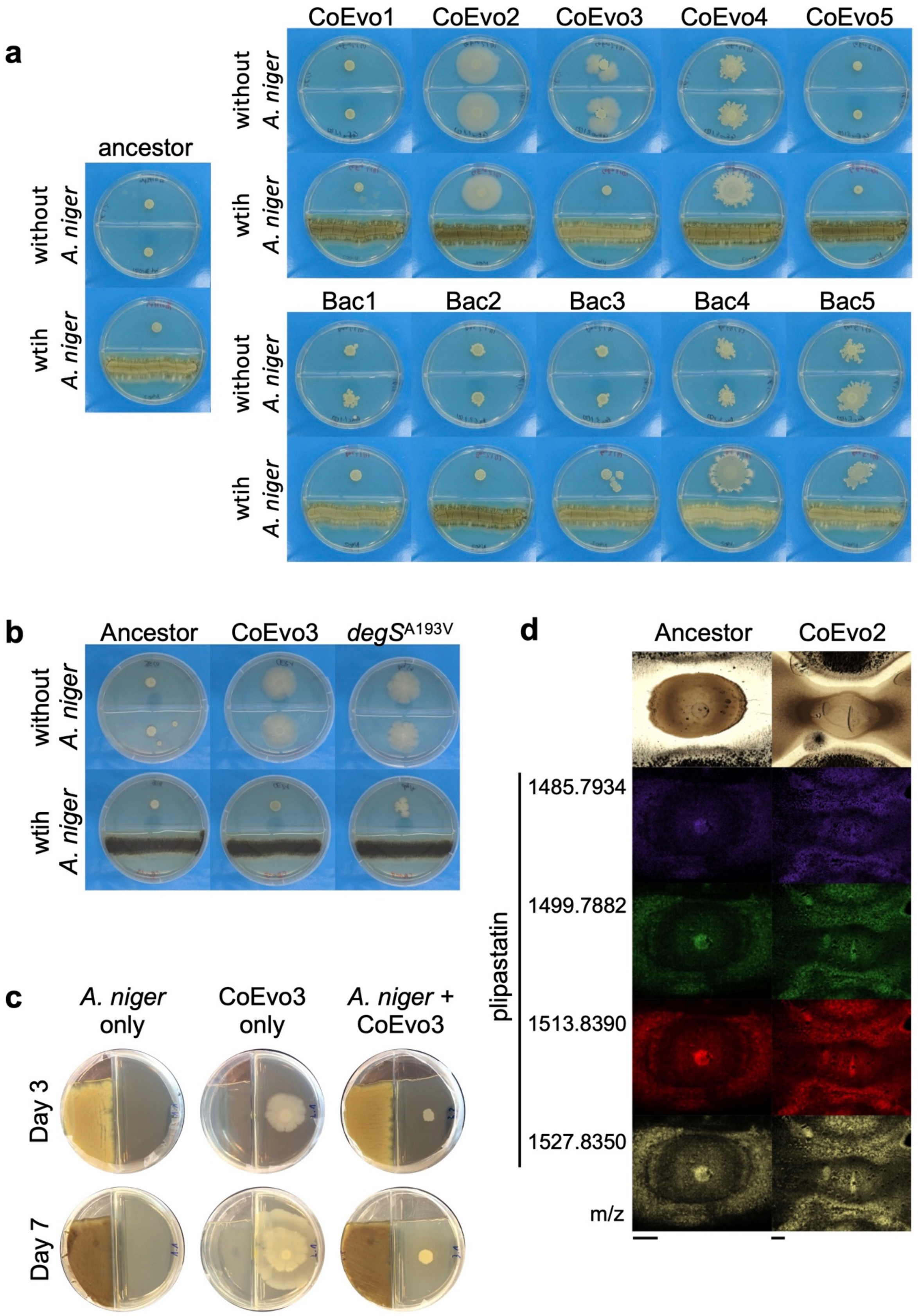


**Fig. S3 | The effects of volatile compounds on *B. subtilis* growth, and spatial detection of plipastatin in CoEvo2. a,** Colony spreading of the ancestor and evolved isolates in the absence (top panels) and presence of *A. niger* (lower panels). **b,** Colony spreading of the ancestor, CoEvo3 and *degS*$^{A193V}$ mutant in the absence (top panels) and presence of *A. niger* (lower panels). **c,** Experimental setup used to trap VOCs at day 3 and 7. The empty space in the agar medium was used to place the steel traps containing 150 mg Tenax TA and 150 mg Carbopack B. **d,** MALDI-MSI spatial detection of plipastatin isoforms in bacterial colonies (wild-type and CoEvo2) grown between two fungal streak lines. m/z values of surfactin isoforms are indicated on the left. Scale bars = 2 mm.

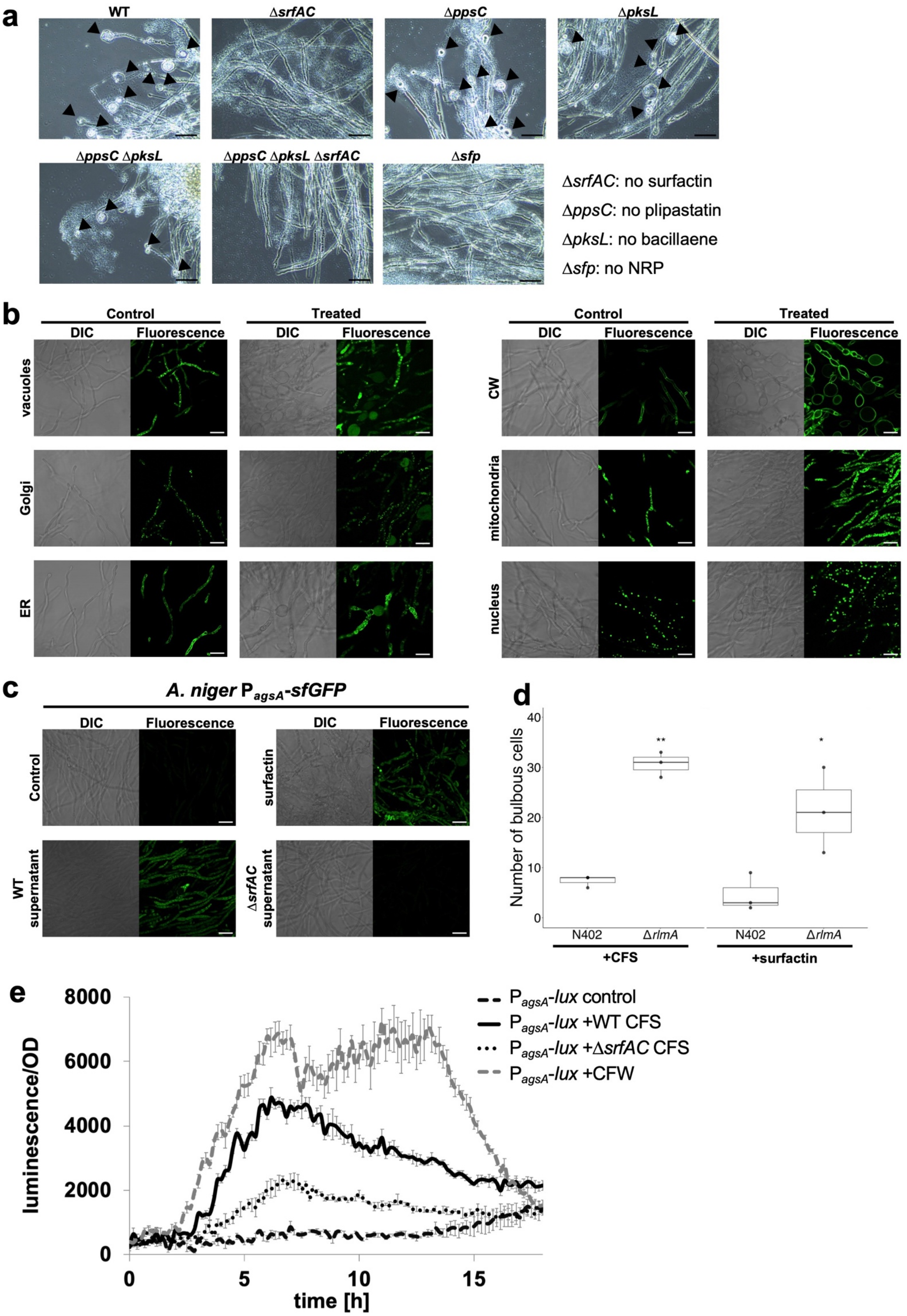
a
WT
ΔsrfAC
ΔppsC
ΔpksL
ΔppsC ΔpksL
ΔppsC ΔpksL ΔsrfAC
Δsfp
ΔsrfAC: no surfactin
ΔppsC: no plipastatin
ΔpksL: no bacillaene
Δsfp: no NRP
b
Control
Treated
DIC
Fluorescence
vacuoles
Golgi
ER
CW
mitochondria
nucleus
c
A. niger PagsA-sfGFP
DIC
Fluorescence
Control
surfactin
WT supernatant
ΔsrfAC supernatant
d
Number of bulbous cells
40
30
20
10
0
**
*
N402
ΔrlmA
N402
ΔrlmA
+CFS
+surfactin
e
PagsA-lux control
PagsA-lux +WT CFS
PagsA-lux +ΔsrfAC CFS
PagsA-lux +CFW
luminescence/OD
8000
6000
4000
2000
0
0
5
10
15
time [h]

**Fig. S4 | Influence of surfactin on fungal hyphae. a,** Microscopy visualisation of bulging fungal hyphae with wild type (WT) and various mutants, including strain lacking surfactin (Δ*srfAC*), plipastatin (Δ*ppsC*), bacillaene (Δ*pksL*), plipastatin and bacillaene (Δ*ppsC* Δ*pksL*), plipastatin, bacillaene, and surfactin (Δ*ppsC* Δ*pksL* Δ*srfAC*), or all non-ribosomal peptides (Δ*sfp*). Scale bar = 20 μm. **b,** DIC (left) and green fluorescence (right) imaging of the *A. niger* MA23.1.1 strain for vacuoles ($P_{gpdA}$-CpyA::eGFP-T*trpC*); Ren1.10 strain for Golgi ($P_{gmtA}$-eYFP:: GMTA-T*gmtA*), MA141.1 strain for endoplasmic reticulum, ER ($P_{gpdA}$-*glaA*:: sGFP-HDEL-T*trpC*); AR0#11 strain for cell wall, CW ($P_{gpdA}$-*glaA*::sGFP-T*trpC*); BN38.9 strain for mitochondria ($P_{gpdA}$-CitA::eGFP-T*trpC*); and MA26.1 strain for nucleus ($P_{gpdA}$-H2B::eGFP-T*trpC*) in the absence (Control) or presence (Treated) of bacterial cell free supernatant. Scale bar = 20 μm. **c,** DIC (left) and green fluorescence (right) imaging of the *A. niger* JvD1.1 strain carrying $P_{agsA}$-eGFP-T*trpC* for detection of *agsA* gene expression in the absence (control) and presence of cell-free WT supernatant, 20 μg/ml surfactin, and cell-free Δ*srfAC* supernatant. Scale bar = 20 μm. **d,** Number of bulbous cells by the wild-type N402 and Δ*rlmA* mutant *A. niger* in the presence of cell-free WT supernatant (CFS) or 20 μg/ml surfactin. Student's t-test with Bonferroni-Holm correction was performed (*$p_{adjust}$ <0.05, **$p_{adjust}$ <0.01). **e,** Luminescence reporter assay using *A. niger* strains MA297.3 containing $P_{agsA(3\times RlmA\ box)}$ before the promoter-less luciferase. Cultures were treated with LB medium (black dashed line), cell-free WT supernatant (CFS, solid line), cell-free Δ*srfAC* supernatant (Δ*srfAC* CSF, dotted line), or Calcofluor White (CFW, grey dashed line).

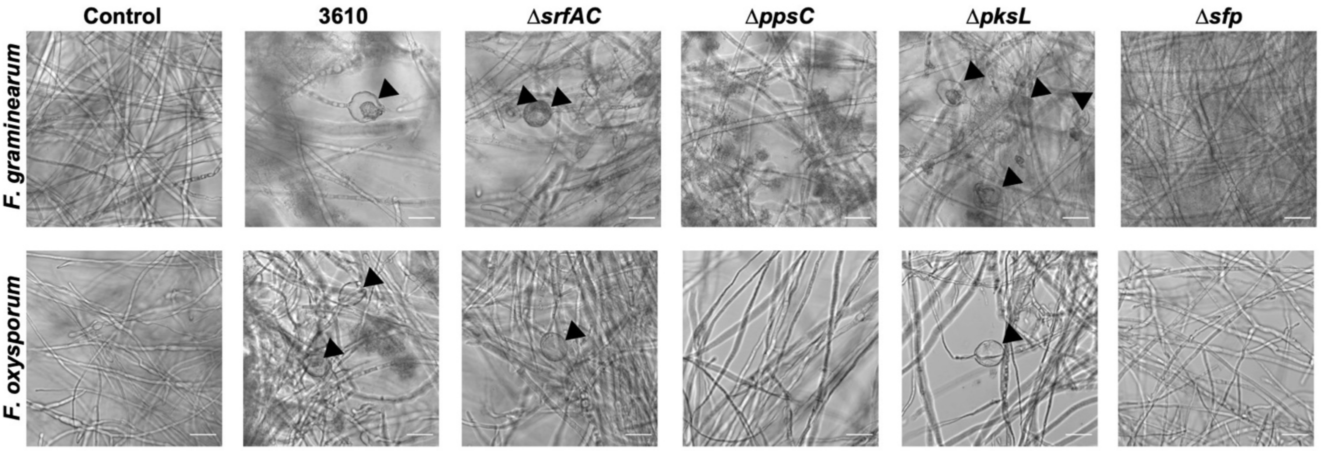


**Fig. S5 | Influence of plipastatin on hyphae of Fusarium species.** Microscopy visualisation of bulging Fusarium hyphae with wild type (WT) and various mutants, including strain lacking surfactin (Δ*srfAC*), plipastatin (Δ*ppsC*), bacillaene (Δ*pksL*), or all non-ribosomal peptides (Δ*sfp*). Scale bar = 25 μm.
