## Supplementary Table 1 for "Enhanced niche colonisation and competition during bacterial adaptation to a fungus"

Supplementary Table 1 for strains, plasmids and oligos

| *B. subtilis* strains | Genotype, description | Reference |
|---|---|---|
| NCIB 3610 | undomesticated wild type strain | [1,2] |
| DK1042 | NCIB 3610, but *comI*$^{Q12L}$ (naturally competent) | [3] |
| 168 *degU* | *trpC* Δ*degU*::Km$^R$ | [4] |
| TB742 | DK1042 Δ*degU*::Km$^R$ | This work |
| TB938 | DK1042 *degU*$^{S202G}$ | This work |
| TB939 | DK1042 *degS*$^{A193V}$ | This work |
| DS4085 | NCIB 3610 Δ*pksL*::Cm$^R$ | [5] |
| DS4114 | NCIB 3610 Δ*ppsC*::Tet$^R$ | [5] |
| DS1122 | NCIB 3610 *srfAC*::Tn10 Spec$^R$ | [6] |
| DS3337 | NCIB 3610 Δ*sfp*::Mls$^R$ | [7] |
| DS4113 | NCIB 3610 Δ*ppsC*::Tet$^R$ Δ*pksL*::Cm$^R$ | [5] |
| DS4124 | NCIB 3610 Δ*ppsC*::Tet$^R$ Δ*pksL*::Cm$^R$ *srfAC*::Tn10 Spec$^R$ | [5] |
| *A. niger* strains | Genotype | Reference |
| N402 | wild-type fungal strain | [8] |
| MA297.3 | N402 $P_{agsA(3\times RlmA\ box)}$-m*luc*-T*trpC*-*pyrG*** | This work |
| MA584.2 | N402 $P_{agsA(RlmA\ box\ mutated)}$-m*luc*-T*trpC*-*pyrG*** | This work |
| Δ*rlmA* | N402 Δ*rlmA*::hyg$^R$ | [9] |
| AR0#11 | N402 $P_{gpdA}$-*glaA*::sGFP-T*trpC* | [10] |
| MA141.1 | N402 $P_{gpdA}$-*glaA*:: sGFP-HDEL-T*trpC* | [11] |
| Ren1.10 | N402 $P_{gmtA}$-eYFP:: GMTA-T*gmtA* | [11] |
| MA23.1.1 | N402 $P_{gpdA}$-CpyA::eGFP-T*trpC* | [12] |
| FG7 | N402 $P_{synA}$-eGFP::SynA-T*synA* | [13] |
| BN38.9 | N402 $P_{gpdA}$-CitA::eGFP-T*trpC* | [14] |
| MA26.1 | N402 $P_{gpdA}$-H2B::eGFP-T*trpC* | [12] |
| JvD1.1 | N402 $P_{agsA}$-eGFP-T*trpC* | [15] |
| Fungal strains | | |
| *A. awamori* | FSU 11418 | JMRC |
| *A. brasiliensis* | FSU 35902 (DSM 1988) | JMRC |
| *A. tubingiensis* | FSU 11408 | JMRC |
| *A. nidulans* | HKI G034 | JMRC |
| *F. graminearum* | IBT 41925 | IBT |
| *F. oxysporum* | IBT 40872 | IBT |
| JMRC: Jena Microbial Resource Collection at Leibniz Institute for Natural Product Research and Infection Biology Hans Knöll Institute, Jena, Germany (https://www.leibniz-hki.de/en/jena-microbial-resource-collection.html)<br>IBT: IBT Culture Collection at DTU Bioengineering, Kongens Lyngby, Denmark (https://www.bioengineering.dtu.dk/research/strain-collections/ibt-culture-collection-of-fungi) | | |
| Plasmids | description | Source |
| pMiniMad | *ori*$^{BsTs}$ *Amp*$^R$ *Mls*$^R$ | [16] |
| pTB693 | pMiniMad with *degU*$^{S202G}$ | This work |
| pTB694 | pMiniMad with *degS*$^{A193V}$ | This work |
| pMA334 | vector with *pyrG* flanking regions | [17] |

| | | | |
|---|---|---|---|
| pBN008 | | vector with $P_{agsA}$(0.55-kb-rlm2add)-*uidA-pyrG* | [9] |
| pVG4.1 | | vector with m*luc*-T*trpC* | [18] |
| pMA348 | | pMA334 with $P_{agsA(3\times \mathrm{RlmA\text{-}box})}$-m*luc*-T*trpC-pyrG*** | This work |
| pMA370 | | pMA334 with $P_{agsA(\mathrm{RlmA\text{-}box\ mutated})}$-m*luc*-T*trpC-pyrG*** | This work |
| Oligos | | sequence | gene targeted |
| oAR23 | *Nco*I | ATCCATGGTGGCGGCTGAGAAGTCGTCG | *degU* in CoEvo2 |
| oAR24 | *Bam*HI | GCGGATCCAAGAGGTTATCTGCTGAAAG | *degU* in CoEvo2 |
| oAR30 | *Sal*I | CCGTCGACTTGGCGATAAACTTGAAGTG | *degS* in CoEvo3 |
| oAR41 | *Bam*HI | ATGGATCCTGAAGAGCGCAACCTCAAAC | *degS* in CoEvo3 |
| oAR25 | | AGACTTGCCAAGCTCTTC | *degU* |
| oAR26 | | GCTTGTAGAGCTGTATCC | *degU* |
| oAR31 | | TCAGGTCGAACCCTTTAC | *degS* |
| oAR32 | | AACAGCTGGTCGAAGAAC | *degS* |
| oAR27 | | TCCTCTGGCCATTGCTCTG | pMiniMad MCS |
| oAR28 | | CGAAGTTAGGCTGGTAAG | pMiniMad MCS |
| PagsAP1f-NotI | | GCGGCCGCTCTAGAACTAGT | |
| TtrpCP2r-NotI | | AAGGAAAAAAGCGGCCGCTCTAGAAAGAAGGATTACCTC | |
| PagsAP4f-NotI | | AAGGAAAAAAGCGGCCGCCTGCAGTAGTGGCGGCTGCTTC | |
| PagsA-AF-R-mut-RlmA2 | | CTCGGTGGTCGCCGCCGAGAAACGTCATATCAGGATAGC | |
| PagsA-AF-F-mut-RlmA1 | | ATATGACGTTTCTCGGCGGCGACCACCGAGAGTAGAGAATGA | |
| PagsAP2r | | CTCGATCTTTCTGCGACCCATGATGGCAAGCGGCGTGTGGTA | |
| oBK7 | | CCGAGTACAAGGARGCCTTC | CaM |
| oBK8 | | CCGATRGAGGTCATRACGTGG | CaM |
